# Phenotype and function of IL-10 producing NK cells in individuals with malaria experience

**DOI:** 10.1101/2024.05.11.593687

**Authors:** Sarah A. McNitt, Jenna K. Dick, Maria Hernandez Castaneda, Jules A. Sangala, Mark Pierson, Marissa Macchietto, Kristina S. Burrack, Peter D. Crompton, Karl B. Seydel, Sara E. Hamilton, Geoffrey T. Hart

## Abstract

*Plasmodium falciparum* infection can trigger high levels of inflammation that lead to fever and sometimes severe disease. People living in malaria-endemic areas gradually develop resistance to symptomatic malaria and control both parasite numbers and the inflammatory response. We previously found that adaptive natural killer (NK) cells correlate with reduced parasite load and protection from symptoms. We also previously found that murine NK cell production of IL-10 can protect mice from experimental cerebral malaria. Human NK cells can also secrete IL-10, but it was unknown what NK cell subsets produce IL-10 and if this is affected by malaria experience. We hypothesize that NK cell immunoregulation may lower inflammation and reduce fever induction. Here, we show that NK cells from subjects with malaria experience make significantly more IL-10 than subjects with no malaria experience. We then determined the proportions of NK cells that are cytotoxic and produce interferon gamma and/or IL-10 and identified a signature of adaptive and checkpoint molecules on IL-10-producing NK cells. Lastly, we find that co-culture with primary monocytes, *Plasmodium*-infected RBCs, and antibody induces IL-10 production by NK cells. These data suggest that NK cells may contribute to protection from malaria symptoms via IL-10 production.

## INTRODUCTION

Malaria is a mosquito-borne infectious disease that claims roughly 600,000 lives per year and is caused by *Plasmodium* parasites [1]. Over 90% of the world’s malaria deaths occur among children living in sub-Saharan Africa. In humans, *Plasmodium* infection starts by mosquitoes taking a blood meal and injecting sporozoites into the skin whereupon the sporozoites traffic to the liver. After 7 days, infected hepatocytes rupture, releasing thousands of merozoites that start the erythrocytic stage by infecting red blood cells. Lastly, some parasites transform into gametocytes; this is the sexual form of *Plasmodium* that can transmit the parasite back to mosquitoes with a subsequent blood meal. Broadly, this process is characterized into three stages: the sporozoite (liver), erythrocytic (blood), and gametocyte (sexual) stages[2]. Recent headway has been made with malaria sporozoite stage vaccines and monoclonal antibody development[3–5]. These approaches target the circumsporozoite protein (CSP) to block infection progression or reduce the infectious dose of merozoites released from the liver. However, for people who progress to the erythrocytic stage of infection, where CSP is not expressed, susceptibility may remain to the symptomatic and severe manifestations of malaria, including cerebral malaria and severe malaria anemia.

During the erythrocytic stage of infection, the optimal host immune response maintains a delicate balance between curbing parasite growth and minimizing host immunopathology. Proposed mechanisms to kill *Plasmodium*-infected red blood cells (RBCs) and limit parasite growth are thought to include phagocytosis and direct cytotoxic killing of infected RBCs[6–12]. Recently, NK cells have been shown to inhibit *Plasmodium falciparum* growth *in vitro* through antibody dependent cellular cytotoxicity (ADCC)[6]. Not all subsets of NK cells are equally adept at ADCC. Previous work showed that trained subsets of NK cells, originally defined in the context of cytomegalovirus (CMV) infection and referred to as adaptive NK cells, display memory-like functions and enhanced ADCC capabilities[13–17]. Characteristic phenotypes associated with adaptive NK cells are *increased* expression of NKG2C and CD57, activation and maturation markers respectively[16, 17], and *reduced* expression of the signaling adapter Fc receptor γ chain (Fcer1g) and transcription factor promyelocytic leukemia zinc-finger protein (PLZF)[14, 18]. We and others have found that individuals with malaria experience have an adaptive NK cell subset that lacks Fc receptor γ chain and PLZF and correlates with reduced parasitemia and protection from malaria symptoms[7, 8].

The adaptive NK cell subset identified in individuals with malaria experience was also enhanced for both cytotoxic and inflammatory cytokine production *in vitro*[7, 8]. This finding, combined with data showing that NK cell ADCC inhibits the growth of *Plasmodium falciparum in vitro,* supports the hypothesis that this adaptive NK cell subset contributes to reducing parasite load *in vivo.* Specifically when parasite load is low, the development of malaria symptoms does not directly correlate with parasite load, as many individuals maintain significant parasite burden while remaining asymptomatic[11, 19, 20]. Considering IFNγ, a strongly pro-inflammatory cytokine, has been shown to be produced by adaptive NK cells, it is somewhat surprising that their increased proportion correlates with reduced symptoms (e.g. fever), given that inflammation is a driver of fever. Therefore, we explored the hypothesis that the role of NK cells during malaria infection may extend beyond direct parasite control and include anti-inflammatory mechanisms that prevent symptomatic malaria.

In malaria-naïve individuals, *Plasmodium* infection leads to robust production of inflammatory cytokines such as TNFα, IL-1β, IL-12, IL-6, and IFNγ and subsequent development of fever. Controlling this response is critical to prevent aberrant inflammation and damage to host tissues[21–23]. As an important component of this regulation, IL-10 suppresses inflammation during malaria and other infections by dampening the production of pro-inflammatory cytokines, downregulating MHC-II on antigen presenting cells[24], and promoting humoral immunity[25–27]. IL-10 is made by many cell types but is most associated with CD4+ T cells[28–32]. Systemic cytokines are detected in both non-severe and severe malaria, and the ratio of IL-10 levels to inflammatory cytokines can be an indicator of disease[33]. We and others have shown NK cells can make IL-10, but it is unknown how much systemic IL-10 is derived from NK cells, and what role NK cell-produced IL-10 plays during malaria infection.

Our data from the experimental cerebral malaria (ECM) mouse model showed that NK cells can be induced to produce IL-10, which protected mice from severe disease[34]. Additionally, we found that secretion of IL-10 by human malaria-naïve NK cells is induced *in vitro* by cytokines IL-15, IL-21, and IL-12. Cytokines IL-15 and IL-21 are key promotors of NK cell survival and proliferation, while IL-12 is an effector cytokine produced by myeloid cells that can stimulate NK cell activation. The IL-10 made by human NK cells was detected via ELISA and real time PCR, but the proportion and phenotype of NK cells that produce IL-10 was unknown.

In this study, we analyzed blood from adolescents and young adults from Mali, Africa—a malaria endemic country—who have, by this age, experienced multiple exposures to *Plasmodium falciparum*. Some of these individuals still develop symptomatic malaria infection, which is defined by having a fever (>37.5°C) and the presence of parasites in the blood (>2500 parasites/μl). We also analyzed samples from individuals living in the United States with no previous exposure to *Plasmodium* species. We found that individuals with malaria experience had a significantly higher proportion of NK cells producing IL-10 relative to malaria-naïve individuals. Some NK cells were enriched for cytolytic activity, IFNγ, and/or IL-10 production, but others performed multiple functions. IL-10 producing NK cells expressed a unique adaptive and checkpoint marker signature. Lastly, when NK cells were incubated with *Plasmodium falciparum-*infected RBCs in the presence of monocytes and anti-*Plasmodium* antibodies, the NK cells were induced to make IL-10. This data set demonstrates that IL-10-producing NK cells are more prevalent in individuals living in this malaria endemic area and suggests that human NK cells producing IL-10 may have a protective role in malaria infection.

## RESULTS

### NK cells from individuals in malaria-endemic regions secrete more IL-10 than NK cells from malaria-naïve individuals

The observation that NK cells could rescue mice from ECM death in a manner that was dependent on IL-10 secretion led us to hypothesize that IL-10 release from NK cells may also play a protective role in human *Plasmodium* infection. We sought to determine whether NK cells from individuals who have undergone many seasonal exposures to *Plasmodium* infections have a higher propensity to secrete IL-10 than those from malaria-naïve individuals. To test this hypothesis *in vitro*, we exposed NK cells from healthy donors that were either malaria-naïve (USA) or had malaria experience (Mali) to inflammatory conditions using a previously published 6-day cytokine stimulation procedure (Figure 1A)[34]. We then measured the release of IL-10 using an extracellular IL-10 capture reagent. In line with our previous work showing that human NK cells secrete minimal IL-10 with IL-15 treatment alone[34], we observed that less than 1% of malaria-naïve and malaria-experienced NK cells release IL-10 under these conditions (Figure 1B-D). Incubation with IL-15, IL-21 and IL-12 led to significantly greater IL-10 secretion from NK cells from both malaria-naïve and malaria-experienced individuals. Surprisingly, about ∼20-fold more NK cells from malaria-experienced individuals secreted IL-10 as compared to malaria-naïve controls (Figure 1C-D). We also examined individuals with malaria experience that were still susceptible to clinical malaria over the course of the malaria season. This sampling occurred before the start of the malaria season (Pre-Malaria), when the person presented with clinical malaria (Malaria), and one week following diagnosis and treatment (Convalescent) (Figure 1E)[11]. Overall, there was a relatively stable frequency of NK cells producing IL-10 before, during, and after an individual presented with clinical malaria.

**Figure 1.**
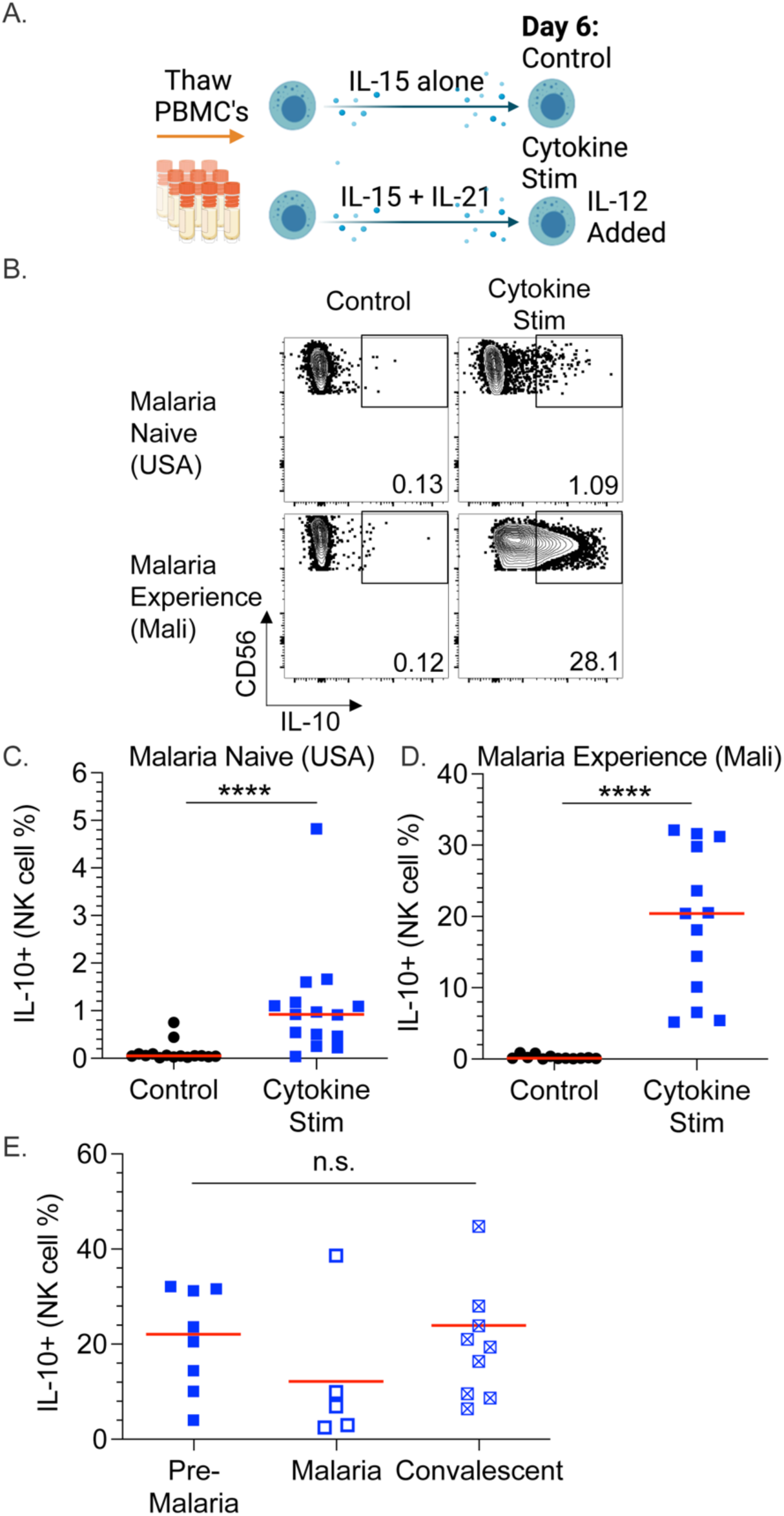
NK cells from malaria-experienced individuals (Mali) secrete more IL-10 than NK cells from malaria-naïve (USA) individuals. (A) Experimental setup for cytokine stimulation. (B) Example flow cytometry staining of IL-10 production by CD56+ NK cells treated with IL-15 alone (Left)(control) or with a combination of IL-15 and IL-21 for 6 days followed by IL-12 on Day 7 (Right)(Cytokine stim). Malaria-naïve (USA)(Top) and malaria-experienced (Mali)(Bottom) individuals. (C-D) Comparison of IL-10 production from malaria-naïve (USA)(C) and malaria-experienced (Mali)(D) individuals. (E) Comparison of IL-10 production from malaria-experienced individuals before the malaria season (Pre-Malaria), when they presented with malaria (Malaria), and 7 days after they were treated for malaria (Convalescent). Red horizonal lines represent median values. ****p<0.0001 as determined by Wilcoxon signed-rank test (C,D) or one-way ANOVA with Tukey’s multiple comparison test (E).

In addition to cytokine stimulation alone, we also assessed IL-10 secretion by NK cells during ADCC and natural cytotoxicity assays in which NK cells become activated through antibody binding to the Fc receptor (FcγRIIIa / CD16a) or through recognition of stress ligands in the absence of human leukocyte antigens (HLA) respectively (natural cytotoxicity) (Figure 2A). Following cytokine stimulation, IL-10 release from NK cells was substantially greater from malaria-experienced individuals in both ADCC (∼10-fold increase) and natural cytotoxicity assays (∼5-fold increase) (Figure 2B-D) and was greatest in the ADCC assay (Figure 2C-D). Because NK cell adaptive phenotype and function can be influenced by prior cytomegalovirus (CMV) exposure, we assessed IL-10 production in CMV positive and negative individuals. Although all malaria-experienced individuals were also cytomegalovirus (CMV) positive, no difference in NK cell IL-10 secretion was observed in USA subjects that were CMV positive or negative during cytokine stimulation, ADCC, or natural cytotoxicity assays (Supplemental Figure 1A). IL-10 production was not different in NK cells sampled from individuals prior to, during, or after malaria in either cytolytic assay (Figure 2E-F). Taken together these data indicate that the NK cells from malaria-experienced individuals more robustly secrete IL-10 compared to those from malaria-naïve individuals. Among the assays tested, exposure to cytokines IL-15, IL-21, and IL-12 plus the addition of an ADCC stimulus causes NK cells to produce the most IL-10.

**Figure 2.**
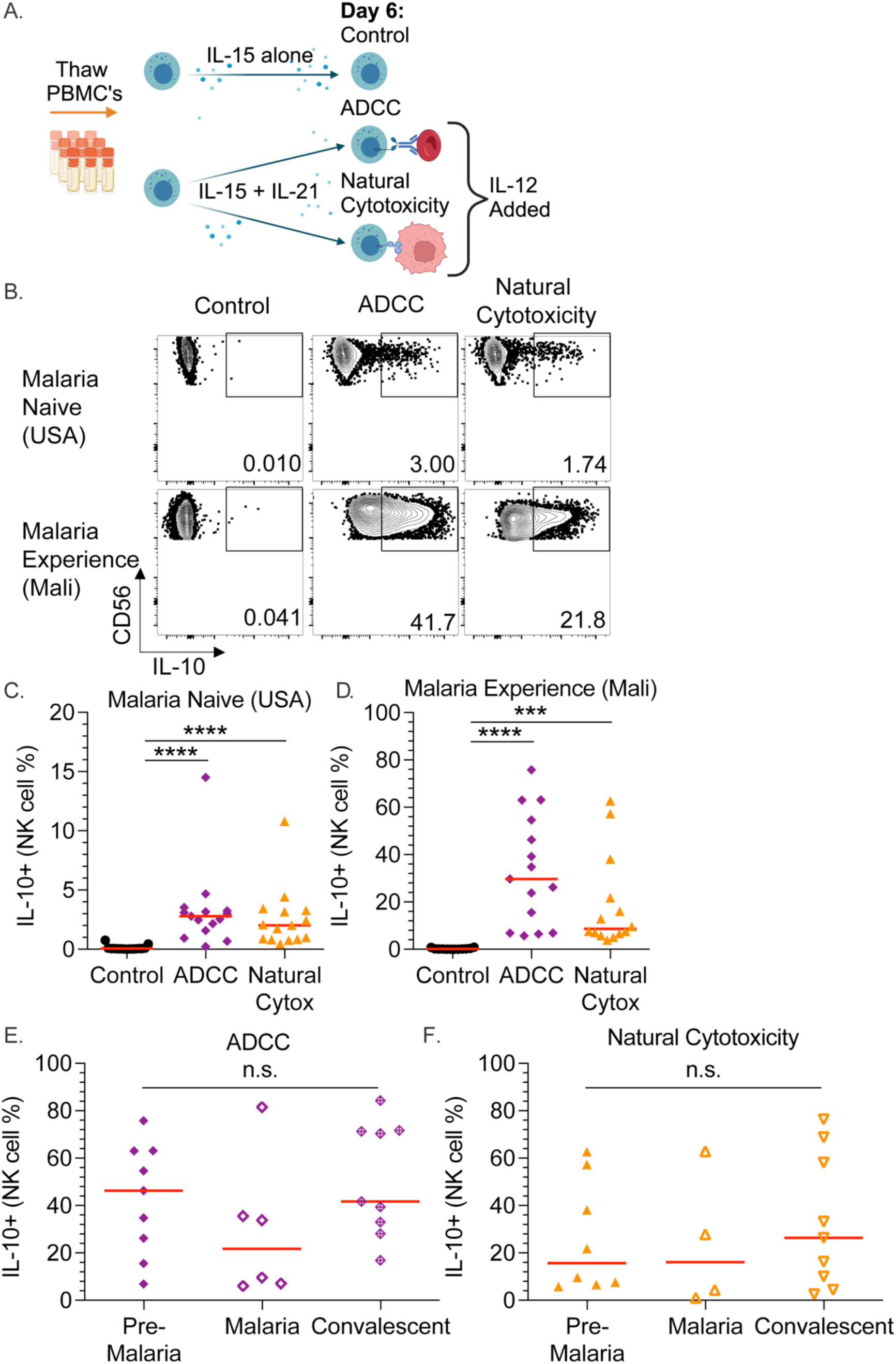
NK cells produce IL-10 during antibody dependent cellular cytotoxicity (ADCC) and natural cytotoxicity. (A) Experimental setup for ADCC and natural cytotoxicity assays. (B) IL-10 production from NK cells treated with IL-15 alone (control)(Left), with IL-15 and IL-21 for 6 days followed by IL-12 and either an ADCC (Middle) or natural cytotoxicity (Right) assay. (C-D) Comparison of ADCC or natural cytotoxicity-induced IL-10 production from the NK cells of malaria-naïve (C) or malaria-experienced (D) individuals. (E-F) Comparison of IL-10 production from malaria susceptible individuals before the malaria season (Pre-Malaria), when they got malaria (Malaria), and 7 days after they were treated for malaria (Convalescent) for ADCC (E) or Natural Cytotoxicity (F). Red horizonal lines represent median values ***p<0.001 ****p<0.001 as determined by one-way ANOVA with Tukey’s multiple comparison test (C-F).

### NK cells co-produce IL-10, IFNγ, and CD107a

From the observations noted above, we postulated that IL-10 may be secreted at the same time as cytotoxic degranulation. When NK cells degranulate their cytolytic granules, the granule membrane fuses with the cell membrane exposing CD107a (LAMP-1) on the surface of the cell. NK cells are also known to produce multiple pro-inflammatory cytokines, such as IFNγ, during degranulation. To assess production of all three molecules, we captured IL-10 release, stained intracellularly for IFNγ, and monitored surface expression of CD107a (gating strategy shown in Supplement Figure 1B). As expected, IFNγ was readily produced by NK cells in all conditions (Figure 3A). CD107a was also detected in all assays but was greatest in the natural cytotoxicity assay (Figure 3A). In a pairwise analysis, co-expression of IFNγ and IL-10 was evident (Figure 3B-D) as well as co-expression of CD107a and IL-10 (Figure 3C-D). Using SPICE (“Simplified Presentation of Incredibly Complex Evaluations”) analysis, we compared the proportions of cells that were expressing CD107a, IFNγ, and IL-10 as single, double, and triple NK cell producers in all three stimulation assays (Figure 3D). Cytokine and ADCC stimulation generated similar patterns of effector molecule expression, with most cells expressing IFNγ, followed by IL-10 and IFNγ co-expression. We also observed some triple producers (IFNγ, IL-10, CD107a) but few cells that expressed only IL-10 or CD107a. During natural cytotoxicity, most cells produced both CD107a and IFNγ. The frequency of cells secreting IL-10 was reduced and was mainly found in conjunction with both IFNγ and CD107a.

**Figure 3.**
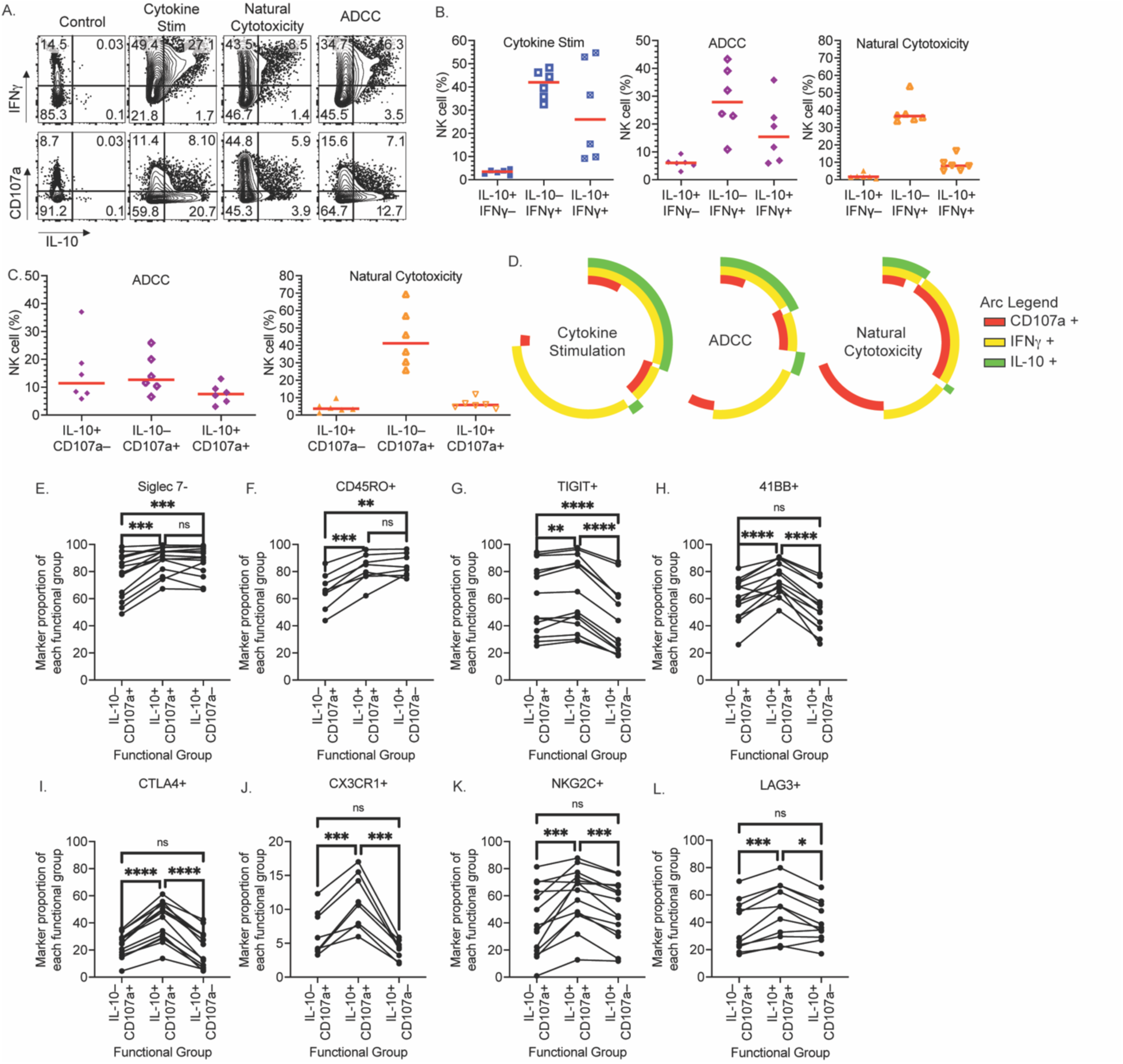
IL-10 is co-expressed with IFNγ and CD107a. (A) Representative flow cytometry data of IL-10 by IFNγ (Top) or by CD107a (Bottom) following IL-15 alone (Control)(Left) or IL-15 and IL-21 for 6 days and then with cytokine stimulation (IL-12)(Middle left), natural cytotoxicity (No IL-12)(Middle right), or ADCC (No IL-12)(Right) assay. (B) For individuals with malaria experience, proportions of Boolean gates of IL-10 and IFNγ for cytokine stimulation (Left), ADCC (Middle), or natural cytotoxicity (Right). (C) For individuals with malaria experience, proportions of Boolean gates of IL-10 and CD107a for ADCC (Left) or natural cytotoxicity (Right). (D) SPICE analysis of the co-expression of IL-10, IFNγ, and CD107a following cytokine stimulation, ADCC, or natural cytotoxicity. (E-L) For an ADCC assay, first, cells of each functional group were gated (selected) being IL-10-/CD107a+, IL-10+/CD107a+, or IL-10+/CD107a–. Then the proportion of each NK cell marker was determined. The proportion of each NK cell marker as a percent of each functional group shown. Red horizonal lines are median. ** = p-value <0.01, *** = p-value <0.001, **** = p-value <0.0001 by a one-way ANOVA with Tukey’s multiple comparison test.

The phenotype of IL-10 secreting NK cells has not been reported. To discover extracellular markers that correlate with IL-10 release in an unbiased manner, we utilized a 350-marker flow cytometry screening panel and single-cell RNA sequencing to analyze cytokine stimulated malaria-naïve NK cells. Results from these screens revealed several markers that were preferentially expressed on IL-10+ NK cells compared to IL-10– NK cells (Supplemental Figure 2A-B). We noted that several of these — CTLA-4, 4-1BB, PD-1, LAG-3 (CD233), Siglec-7, KLRG1, and TIGIT — have been characterized as immune checkpoint molecules in NK cells[35–40]. We also identified markers less studied in NK cell biology such as CX3CR1, CD45RO, and CRTAM[41, 42]. These markers were integrated into the study’s flow cytometry assays as well as markers associated with adaptive NK cells and viral infection, such as NKG2C, FcRγ chain (Fcer1g), PLZF and CD8[7, 14, 16–18, 43, 44]. Recently, another CD56 negative (neg), CD7+, FcRγ chain negative NK cell population was found to be associated with malaria exposure and protection from malaria, similar to CD56 dim (positive), FcRγ chain negative NK cells[7, 45]. We assessed the proportions of CD56 positive and CD56 neg (CD7+, CD3–) NK cells before (Day 0) and after 6 days of cytokine incubation[8]. We found that after cell culture the CD56 neg NK cell proportion was drastically lowered indicating either death or conversion to a CD56 positive phenotype (Supplemental Figure 2C). Therefore, we chose to assess CD56 positive NK cells for expression of IL-10 secretion from malaria-naïve and malaria-experienced individuals.

In the ADCC assay, which causes the strongest IL-10 production, we found three populations of cells: those that are solely producing IL-10 and are not cytotoxic (IL-10+, CD107a–), those that are only cytotoxic (IL-10–, CD107a+), and those that co-express IL-10 and CD107a (IL-10+, CD107a+). Incorporating the markers we identified in the RNA sequencing and protein screens, we associated markers that were enriched in these functional groups. Siglec-7– and CD45RO+ NK cells were enriched in the IL-10 producers (both single IL-10+, CD107a– and double IL-10+, CD107a+ NK cells) (Figure 3 E-F). TIGIT+ NK cells were enriched in the cytotoxic NK cells (both single IL-10–, CD107a+ and double IL-10+, CD107a+ NK cells) (Figure 3G). Lastly, 4-1BB+, CTLA-4+, CX3CR1+, NKG2C+, and LAG-3+ NK cells were enriched in the dually cytotoxic and IL-10 producing NK cells (Figure 3I-L).

We also noted that stimulation in ADCC or natural cytotoxicity assays directly *ex-vivo* (Day 0) induces IFNγ production in 5-20% of the NK cells (Supplementary Figure 3A-B), whereas there is very little IL-10 made. Instead, it took cytokine stimulation to induce the NK cells to produce IL-10 (Day 6)(Supplementary Fig. 3A-B). This is a similar trend to reports of NK cell IFNγ and IL-10 production *in vivo,* which indicate that IL-10 initiates after an early IFNγ response to infection[34, 46].

### NK cells secreting IL-10 express adaptive and immune checkpoint molecules

We found that PD-1+, 4-1BB+, LAG-3+, KLRG1+, PLZF–, TIGIT– and Siglec-7– NK cells were increased in individuals with malaria experience (Figure 4A). Examining malaria-experienced individuals for common markers across all assays, 4-1BB+, CTLA-4+, CX3CR1+, NKG2C+, TIM-3+, and Siglec-7– NK cells were enriched for IL-10 production (Figure 4B-D). This indicates that NK cells expressing these checkpoint (*e.g.* CTLA-4+, 4-1BB+, TIM-3+, Siglec-7–), adaptive (*e.g.* NKG2C+, Siglec-7–), and trafficking (CX3CR1+) markers are more likely to secrete IL-10. For the ADCC assay, additional markers, including CD16a+, KLRG1+, PD-1+, and TIGIT+, on NK cells were significantly enriched for IL-10 production (Figure 4C). Similar trends were seen in malaria-naïve individuals, although the frequency of NK cells expressing individual molecules did vary significantly (Supplemental Figure 4A). Other markers tested were not significantly enriched for IL-10 on malaria-experienced individuals (Supplemental Figure 4B). We did not uncover a sole marker that predicts IL-10 production by NK cells. However, when assessing the co-expression of LAG-3+, NKG2C+, Siglec-7−, and TIGIT+ NK cells by SPICE analysis (Figure 4E), we found that around 25% of IL-10 secreting NK cells expressed all 4 markers, and greater than 50% of the IL-10 secreting cells expressed at least 3 of the 4.

**Figure 4.**
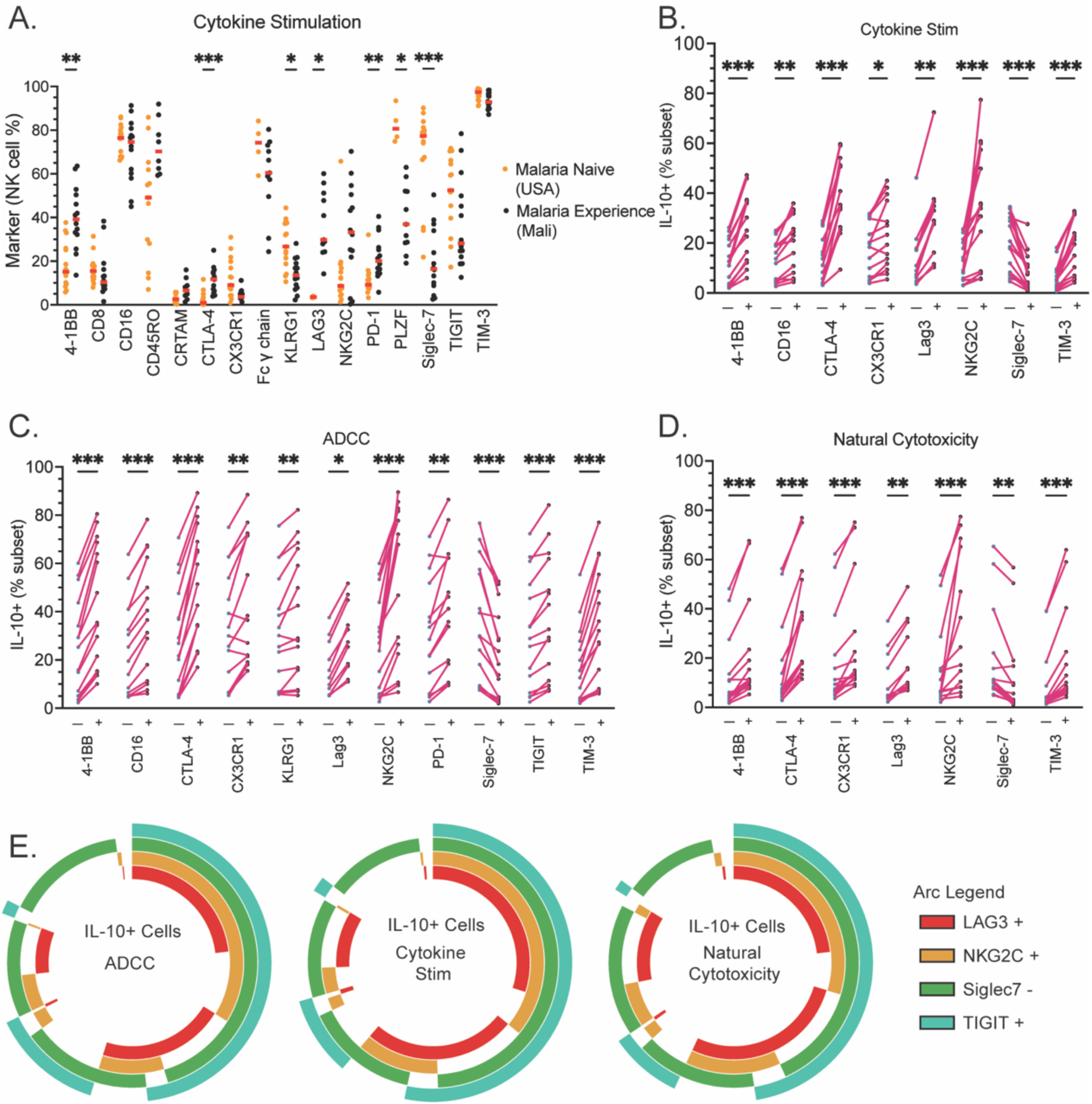
IL-10 production from NK cell subsets for different functional assays for individuals with malaria experience. (A) Proportion of CD56+ NK cells expressing individual markers for malaria-naïve (USA) and malaria-experienced (Mali) individuals after 6 days in IL-15 and IL-21 and stimulation with IL-12 (Cytokine stimulation). (B-D) For individuals with malaria experience, proportion of IL-10 production from NK cell subsets for Cytokine stimulation (B), ADCC (C), and Natural Cytotoxicity (D). SPICE analysis for co-expression of selected markers for all three assays. Data was analyzed using Mann-Whitney non-parametric tests with Bonferroni’s correction (14). * = p-value <0.05, ** = p-value <0.01, *** = p-value <0.001, **** = p-value <0.0001 post Bonferroni correction.

### NK cells produce IL-10 when co-cultured with monocytes, *Plasmodium* infected RBCs, and antibody

Co-culture of NK cells with monocytes and infected RBCs can induce NK cells to produce IFNγ [47]. In the cytokine stimulation assays used in previous experiments, a key component to induce NK cells to produce maximal IL-10 was IL-12. IL-12 is secreted by monocytes and macrophages during NK cell and myeloid cell cross-talk[48]. Monocytes make IL-12 in the process of engulfing infected RBCs[12]. There are many inflammatory molecules in *Plasmodium*-infected RBCs, such as hemozoin, which activates the inflammasome pathway to induce IL-12 and IL-18[49]. Therefore, we hypothesized that during co-culture of NK cells with infected RBCs and monocytes, the IL-12 made by monocytes would stimulate NK cells to produce IL-10. Furthermore, we hypothesized that the addition of anti-RBC antibodies would induce ADCC and potentially further increase IL-10 production. To test this, we used the co-culture conditions laid out in Figure 5. Infected and opsonized RBCs we predict would cause inflammation through hemozoin and an ADCC/ and antibody dependent cellular phagocytosis (ADCP) signals. For the ADCC/ADCP signals, three conditions were used: anti-RBC antibody (αRBC) that binds to both RBCs and infected RBCs, malaria-experienced immune plasma that will bind only infected RBCs, and malaria-naïve plasma that should bind neither RBCs nor infected RBCs. NK cells from malaria-naïve individuals incubated in IL-15 alone for 6 days were used as a negative control for IL-10 production. NK cells from malaria-naïve individuals were also incubated with IL-15 and IL-21 for 6 days, where the proportion of IL-10 production is approximately 2% (Figure 5C). This control was used to make paired fold change comparisons for CD107a, IFNγ, and IL-10 production (Figure 5 A-C). Uninfected RBCs did not cause inflammation but could induce cytokine secretion from NK cells and monocytes if they were opsonized by an anti-RBC antibody causing ADCC and ADCP, respectively. We found that NK cells significantly increased degranulation (CD107a+) when there was an ADCC signal, and NK cells did not degranulate above baseline with infected-RBCs alone (Figure 5A). This is consistent with past evidence showing no significant natural cytotoxicity against infected RBCs[6–8]. In agreement with previous studies, we found that NK cells, when co-cultured with monocytes and infected RBCs, made significantly more IFNγ (Figure 5B)[47, 50]. We also found the NK cells made significant amounts of both IFNγ and IL-10 with the addition of an ADCC/ADCP signal (Figure 5B-C). The magnitude of the IL-10 production was largest in the group that included monocyte, infected-RBCs, and an ADCC/ADCP signal (Figure 5C). This group also had the largest proportion of NK cells that were co-expressing IFNγ and IL-10 (Figure 5D). We conclude that monocytes aide in inducing IL-10 secretion by NK cells and may provide a source of IL-12.

**Figure 5.**
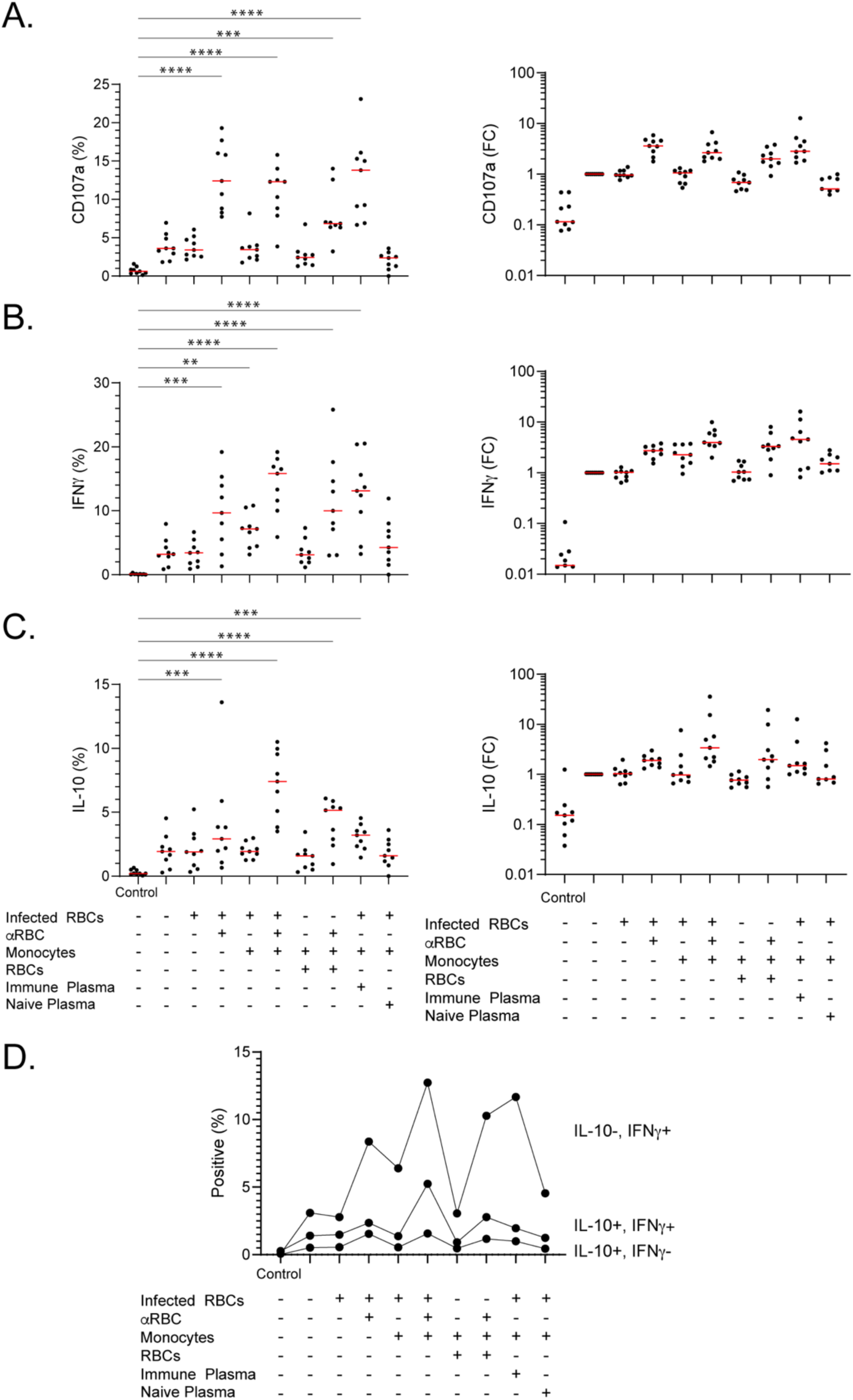
Malaria-naïve NK cell degranulation (CD107a), IFNγ, and IL-10 production in an *in vitro* ADCC/ADCP assay with RBCs or *Plasmodium f.* infected RBCs and purified malaria-naïve monocytes. Column 1 is the Control condition with IL-15 alone. Column 2 is with IL-15 and IL-21. Subsequent conditions have the components on the left column added, indicated by a + symbol. (A-C) Degranulation (CD107a+)(A), IFNγ+ (B), and IL-10+ (C) proportion between all the groups (Left). The fold change (FC) of CD107a+ proportion relative to the IL-15 + IL-21 for 6 days group (Right). Data were analyzed via one-way ANOVA with Tukey’s multiple comparison test done comparing all groups to Control IL-15 alone only group. (D) Comparison across groups of Boolean gates for IL-10 and IFNγ. Red horizontal line is the median. *** = p-value <0.001 **** = p-value <0.0001.

## DISCUSSION

Clinical immunity to *Plasmodium* infection is achieved by controlling both parasite load and inflammation. One mechanism to clear infected RBCs and reduce parasitemia is through cell mediated clearance via Fc receptor recognition of antibodies bound to the surface of infected RBCs. This clearance can be through either phagocytosis by myeloid cells or by NK cell-directed ADCC[6–8, 12, 51–54]. Our previous work demonstrated that adaptive NK cells correlate with reduced parasite load and reduced susceptibility to symptoms, indicating a potential mechanism of acquired trained immunity to malaria. Here, we provide potential insight into the contribution of NK cells to controlling inflammation through the production of the anti-inflammatory cytokine IL-10.

We found that many more NK cells from individuals with malaria infection experience could secrete IL-10 as compared to NK cells from malaria-naive individuals. We hypothesize that this may be important for developing clinical immunity to malaria and preventing severe disease. Although the subjects have undoubtedly been exposed to other infections aside from malaria, it stands to reason that an immune mechanism involved in the control of inflammation would be elevated in individuals experiencing repeated cycles of *Plasmodium* infection in the absence of severe disease. In support of this concept, we have also shown that IL-10 produced by NK cells in a mouse model of experimental cerebral malaria is sufficient to protect from lethality[34]. Additional studies examining NK cells from children who experience severe disease would be useful to determine if the IL-10-producing NK cell population is lacking in these susceptible individuals. Likewise, samples from individuals that experience high or low *Plasmodium* exposure would further strengthen the assertion that NK cells playing a regulatory role through production of IL-10 develop following repeated *Plasmodium* exposure[55]. Previous studies have shown that CD4+ T cells from individuals in malaria endemic areas also display a marked increase in IL-10, which was dependent on malaria infection experience[30]. Thus, there may be multiple types of immune cells engaged to limit inflammation while still allowing for the control of parasite load.

Additionally, we found that most IL-10-producing NK cells also make IFNγ. Production of IFNγ occurred rapidly *ex vivo*; however, NK cells only make IL-10 after being stimulated *in vitro* for 6 days (Supplemental Figure 3). We hypothesize this kinetic difference may be because the immune system first employs NK cells to stimulate the immune response to infection (through early IFNγ), but then uses NK cells to diminish the response via IL-10 secretion and return to homeostasis, thereby preventing overt pathology. Similar ‘waves’ of cytokine production by NK cells have been shown in mouse models of bacterial infection [46]. Here, at the end of the 6-day culture period, we found that human NK cells were capable of concurrently producing IL-10 and IFNγ. This was also observed in CD4+ T cells that were FoxP3– in malaria-experienced children[30]. How these two seemingly contradictory signals would be perceived by responding immune cells is unclear and highlights the importance of understanding how complex cytokine networks impact the outcome of malaria or other infections.

We also found that NK cells could both degranulate (as demonstrated by CD107a+ expression) and secrete IL-10. NK cells have been shown to regulate the contraction phase of immune responses following viral infection through cytotoxic activity against immune cells[56–58]. In conjunction with this activity, IL-10 may also dampen immune responses by inhibiting proliferation, antigen presentation, and the activity of cytokine signals. In contrast to this inhibitory activity, it has also been shown that IL-10 can increase the killing capacity of NK cells[59, 60]. Others have shown that IL-10 induces metabolic changes in NK cells that enhance their cytotoxic functions[59, 60]. From this, we postulated that NK cells secreting IL-10 may increase NK cell degranulation during ADCC and natural cytotoxicity.

Based on dual staining for degranulation and IL-10 production, we also found NK cells that were only degranulating (CD107a+) or only producing IL-10. We found that degranulating cells were enriched for TIGIT+ expression (IL-10+/-) while cells making IL-10 (CD107a+/-) were enriched in the Siglec-7− and CD45RO+ NK cell populations (Figure 3). These markers may identify two subsets of NK cells that have increased function and are also specialized for cytotoxicity or IL-10 production respectively. Siglec-7 is an inhibitory sialic acid binding protein so its loss would theoretically increase functions. TIGIT is also an inhibitory checkpoint molecule so its correlation with increased NK cell degranulation invokes a cell specific and context dependent role for TIGIT in NK cell regulation. There is precedent for checkpoint inhibitors inducing different outcomes when comparing NK cells and T cells. Tim-3+ T cells exhibit poor function, whereas Tim-3+ NK cells exhibit improved function[61, 62]. Our data supports this outcome in NK cells. How and why these checkpoint inhibitors have differing roles in individual cell types has yet to be determined.

Comparing NK cell IL-10 release in response to 3 different stimulations (cytokine stimulation, ADCC, and natural cytotoxicity) showed that higher levels of IL-10 were secreted during ADCC compared to cytokine stimulation alone (Figures 1C and 2C). During ADCC, cytotoxic degranulation leads to the lysis of target cells with subsequent spilling of intracellular components that can promote inflammation[63]. The lysis of infected RBCs has been shown to release molecules, such as heme and hemozoin, that can act as danger associated molecular patterns (DAMPs), which activate multiple inflammatory pathways including toll-like receptor signaling, neutrophil extracellular trap release, and inflammasome formation[64, 65]. Perhaps to dampen this effect, NK cells secrete the anti-inflammatory cytokine IL-10 during or after cytotoxic degranulation. Mechanistic studies into the secretion of IL-10 during NK cell cytotoxic degranulation have not been reported. It is unclear why ADCC provokes more NK cell IL-10 secretion than natural cytotoxicity, but it may be due to differences in signaling pathways. Although there is significant overlap between downstream signaling events, K562 cells—which are used to elicit natural cytotoxicity—primarily express ligands for NKG2D and NCRs, while ADCC signals through CD16a[66].

The finding that IL-10 secretion is only stimulated by IL-15 in the presence of other cytokines, namely IL-21 and IL-12, is consistent with our previous study[34]. IL-12 and IL-18, which are produced by myeloid cells, have effects on NK cell differentiation and can produce memory-like NK cells. Importantly, in addition to secreting IL-12, monocytes have been shown to induce NK cells to produce IFNγ in the presence of infected red blood cells[47, 50]. Our data supports this finding, but also shows that NK cells can make IL-10 under this condition and make even more IL-10 when stimulated to perform ADCC (Figure 5C). Our data indicate that either cell-cell contact or a secreted factor from monocytes induces IL-10 production by NK cells. We hypothesize that IL-12 is important for this effect based on our cytokine stimulation assay, which required the addition of IL-12 for maximal IL-10 production. However, further work will need to be done to determine if cytokines, cell-cell contact, or both are needed to induce the IL-10 production by NK cells.

Lastly, IL-10 production by NK cells across all *in vitro* assays showed that 4-1BB+, CTLA–4+, Siglec-7−, NKG2C+, CX3CR1+, LAG-3+, and TIM-3+ NK cells had significant increases in IL-10 secretion (Figure 4B-D). Additional markers that were significantly increased in the ADCC assay included TIGIT+, KLRG1+, and PD-1+ (Figure 4C). We found a wide range of phenotypic markers on IL-10 producing cells, suggesting that many NK cells can produce this cytokine under inflammatory conditions. This data indicates an enriched signature for a seemingly mixed array of adaptive and exhaustion markers. It also suggests that exhaustion markers are context-dependent and may be beneficial for NK cells by promoting their regulatory functions under certain circumstances (*e.g.* during acute and chronic infections).

Greater IL-10 secretion from the NK cells of malaria-exposed individuals along with evidence of enhanced cytotoxicity and activation of these cells imply a dual role for NK cells as both important for parasite clearance and immunoregulation. NK cells may be key in mediating host immune responses to malaria infection to achieve the needed balance between a response strong enough to clear parasites, but not so forceful as to cause immunopathology.

## MATERIALS AND METHODS

### Human Subjects

Study participants from Kalifabougou, Mali were enrolled in this study as part of an ongoing cohort study of acquired immunity to malaria[11]. Blood was collected by venipuncture for up to 3 different time points for each study participant. The first blood collection was taken during May, before the malaria season began. If participants presented to the local health clinic with malaria symptoms, a second and third blood collection was obtained. The first was obtained during the malaria clinic visit and the second as a convalescent timepoint 7 days later. Blood smears and rapid diagnostic tests (RDT) were performed for malaria parasitemia and diagnosis respectively. Any subject with an RDT+ test was treated for malaria according to the National Malaria Control Program guidelines in Mali, which recommend artemether-lumefantrine for uncomplicated *P. falciparum* malaria. For this study, malaria was qualified as >2500 parasites per μl and a fever above 37.5°C. PBMCs were frozen as described previously[7]. Briefly, after spinning BD Vacutainer® CPT™ Mononuclear Cell Preparation Tube at manufacturer recommended speed(g) and time, plasma was aliquoted and frozen at -80°C. PBMCs were aspirated off the top of the gel, and then the gel was washed once more with RPMI+10%FBS. PBMCs were then counted. PBMCs were resuspended first in solution A (RPMI+50% FBS) in 0.5ml and put in prelabeled cryogenic tubes. Once there were enough samples to freeze, solution B was added (FBS+15%DMSO). Then the tubes were quickly put in a pre-chilled (4°C) “Mr. Frosty” and put in the -80°C freezer for 24 hours. After 24 hours, they were moved to longer term storage in −196°C liquid nitrogen. Samples from Mali were shipped on dry ice (−78.5°C) by courier service to Rockville, MD, where they were again stored in liquid nitrogen. The samples were then forwarded, again on dry ice (−78.5°C) overnight using a courier service to Minneapolis, MN where they were stored in liquid nitrogen before being used for experiments. Malaria-naïve control samples were obtained from Memorial Blood Bank (St. Paul, MN) and PBMCs were processed and frozen as above, except percoll salt gradients (GE; Supplementary Table 1) were used to isolate PBMCs.

### Sex as a biological variable

Both male and female subjects were analyzed in this study, and similar findings are reported for both sexes.

### K562 cell maintenance

K562-articifialAPCs (aAPCs) were maintained between 1x10^5^ to 1.5x10^6^ cells/mL in RPMI-1640 (Fisher, SH3002701) + 10% FBS (PEAK Serum, PS-FB1) (RP10) + 1 μg/mL gentamicin (Sigma-Aldrich, G1272) and incubated at 37 °C with 5% CO_2_. Unless listed otherwise, all 37 °C incubations are conducted at 5% CO_2_.

### Cytokine Stimulation and Cytotoxicity Assays

PBMC vials were thawed, washed, and resuspended in 2 mL of culturing media (Lonza X-VIVO-15, Serum-free hematopoietic cell medium, with L-Glutamine, gentamicin, and phenol red, and 20% heat inactivated human AB serum (Peak Serum)). Culture media was then added to each sample to create a concentration of 800,000 cells per mL. For each sample, 1 mL of this cell suspension was plated for day 0 experiments. Cells were rested overnight at 37°C, 5% CO_2_ before proceeding with cytotoxicity assays and cell staining as described below. To the remaining cell suspension, IL-15 (National Cancer Institute) was added to a final concentration of 20ng/mL. This cell suspension was thoroughly mixed and plated into a 24 well plate to serve as the control day 6 sample. To the remaining cell suspension, recombinant human IL-21 (Biolegend) was added to a final concentration of 50 ng/mL. This cell suspension was then plated at 800,000 cells per well and incubated at 37°C, 5% CO_2_ for 6 days.

For the day 0 functional assays, 500,000 cells were plated into 3 wells (one for the natural cytotoxicity assay, one for ADCC and one for a negative control) per PBMC sample in a 96 well plate. For the ADCC and natural cytotoxicity assays, red blood cells (Memorial blood bank) and K562 cells were resuspended to a concentration of 500,000 cells per 100µL. RBCs were purified from whole blood by leukocyte reduction filtration (Fenwal Inc.) and resuspended at 50% hematocrit in RPMI-1640 and 25 mM HEPES, L-Glutamine and 50 mg/l Hypoxanthine (K-D Medical). For the natural cytotoxicity assay, K562 cells were added to sample PBMC’s at a ratio of 1:1. For the ADCC assays, the RBC suspension was incubated with rabbit anti-human RBC antibodies for 20 minutes (Rockland), then added to 500,000 PBMC’s at a 1:1 ratio in the 96 well plate. For the negative control, 100 µL of culturing media was added. All combinations were incubated at 37°C, 5% CO_2_ for 4 hours. Following the incubation period, a master mix of anti-human CD107a (Biolegend) and IL-10 catch reagent (Miltenyi Biotec) were added per manufacturer’s instruction and cells were incubated for 1 more hour at 37°C in 5% CO_2_.

Day 6 functional assays followed the protocol above with the following exceptions. On day 6, IL-12 was added to all wells (3ng/mL) except those serving as the control (IL-15 only). The cytokine stimulation assay includes PBMC’s stimulated with IL-15 (20ng/mL), IL-21 (50ng/mL) for 6 days and IL-12 (3 ng/mL) for 5 hours. The ADCC and natural cytotoxicity assays were performed as on day 0 with the exception that IL-12 (3ng/mL) was included. Cytokine concentrations were the same for the functional assays as for the cytokine stimulation assay. Quality control of staining was monitored using the same internal control (PBMCs derived from a single donor) in every batch of samples analyzed. For experiments where both IL-10 and IFNγ were stained in the same sample we found that adding IL-12 to the sample induced most of the NK cells to produce IFNγ (data not shown). Therefore, when we did ADCC or natural cytotoxicity assays and assessed IFNγ, we did not add IL-12 (i.e. in Figure 3). We added anti-CD107a, and IL-10 catch reagent into the wells at the beginning of the ADCC and natural cytotoxicity functional tests. Then after 3 hrs., we added brefeldin A and monensin (Sigma) (10μl total volume)(1μg/ml Brefeldin A and monensin final). This allows the IL-10 to be released from the NK cells and detected on the outside of the cell, and then later for IFNγ to be sequestered for detection by internal antibody staining.

### Co-culture NK ADCC and Monocyte Antibody Dependent Cellular Phagocytosis (ADCP) Assay

NK cells were thawed and plated in a 24-well plate at a density of 800,000 cells/well in 1ml Lonza media with IL-15 and IL-21 and incubated in a 37^0^C, 5%CO_2_ incubator for 6 days. On day 6, NK cells were resuspended at 100,000 cells/well and transferred to a 96-well plate. Monocytes were previously purified from whole blood (StemCell Co.) and frozen similarly to PBMCs above. Monocytes were added at 100,000 cells/well, RBCs and *Plasmodium*-infected RBCs (iRBCs) were added at 200,000 cells/well. Mali plasma and US plasma were added at a dilution of 1:10 final concentration. The plate was then incubated for 5 hrs. The plate was spun down at 550xg for 4 minutes, and 115μl of supernatant was carefully removed. 15μl of IL-10 catch reagent (Miltenyi), anti-CD107a (1:200 final) mix was added to each well and incubated for another 2 hrs. Then, after two more hours 10μl of Monensin/BFA solution (1μg/ml final) was added to each well and incubated for two more hours. The cells were then centrifuged, washed, and stained for flow cytometry.

### Flow Cytometry

Cells were resuspended in a master mix composed of PBS and viability dye (Tonbo) and incubated at 4^0^C for 30min. At the end of the 30 (min) incubation, cells were washed with FACS buffer (PBS, 2% FBS, 2 mM EDTA), stained with surface antibodies, and incubated at room temperature in the dark for 20 min. For assays with no internal stain, cells were incubated with 2% formaldehyde for 10 minutes, then washed and resuspended in PBS for flow cytometry analysis. If internal staining was performed, the cells were washed and incubated in 2% formaldehyde (Thermo Fisher Scientific) at 37°C for 10 min. After incubation, the cells were washed and resuspended in 0.04% Triton X-100 (Fisher Scientific) at room temperature in the dark for exactly 7 mins. The cells were then washed and stained with an internal master mix in FACS buffer + 2% BSA. Cells were then incubated at room temperature in the dark for 120 mins. After incubation, the cells were washed once with FACS buffer and resuspended for flow cytometry. Samples were run on a BD Fortessa flow cytometer according to the manufacturer’s instructions and analyzed using FlowJo 10.8.1. SPICE analysis was done using a combination of the Pestle program (version 2.0) and SPICE program (Version 6.1).

### Single cell RNA seq analysis

Single-cell RNA-Seq analysis was carried out using the ddSEQTM Single-Cell Isolator (Bio-Rad, Hercules, CA) and the SureCellTM WTA 3’ Library Prep Kit (Illumina, San Diego, CA). Magnetically enriched NK cells from malaria-naïve subjects were treated with cytokines IL-15, IL-21, IL-12 or IL-15 alone prior to capture on the ddSeq Single-Cell Isolator. Library preparation was done using SureCellTM WTA 3’ Library Prep Kit. Resulting libraries were analyzed for an appropriate length distribution using an Agilent TapeStation, and initially sequenced on the Illumina MiSeq Nano with read lengths: read 1 = 70 bp, index = 8 bp, R2 = 76 bp. Deeper sequencing was then achieved using an Illumina NextSeq 550 High-ouput 2x75 flow cell (total = 6 samples). Per-cell gene expression was quantified for each sample using the BaseSpace® SureCellTM RNA Single-Cell Analysis Workflow v1.2.0 from Illumina, which uses Isas v1.2.6-000184develop for analysis, STAR v2.5.2b for read alignment to the Homo sapiens UCSC hg38 genome, and SAMtools v 1.3. Data was processed downstream using the Seurat single-cell analysis pipeline v4 in R v 4.1.0[67]. Samples were filtered of low quality (number gene Features < 200) cells before normalization with SCTransform[68] using 3,000 variable genes, and batch correction using the Seurat integration procedure using the top 30 PCs[69]. UMAP visualizations and nearest neighbor clustering (Louvain algorithm) were performed using the top 6 PCs. Purified NK cells were used and single cells that were determined CD3 negative by gene expression were 0 and were labeled as IL10 positive if IL10 raw counts ≥ 1. NK cells that were CD3 negative and IL10 negative were compared to cells that were NK cells that were CD3 neg and IL10 positive (baseline group) with the Wilcoxon rank sum test using their SCT-normalized gene counts (assay=“SCT”, min.pct=0.1, logfc.threshold=0, only.pos=FALSE). Genes were considered differentially expressed if FDR < 0.05. Those DEGs were plotted using the Seurat DotPlot() function.

### 350 Surface Protein Analysis (LEGENDScreen^TM^)

To assess the expression of approximately 350 surface proteins at once on NK cells that produced IL-10, cells were analyzed using Biolegend’s LEGENDScreen*^TM^* Human PE Kit (Biolegend, 700007). Seven subjects were incubated with IL-15 and IL-21 (as above) for 6 days. On the 6^th^ day, IL-12 was added to promote NK cells to produce IL-10. The seven subjects were then barcoded with combinations of CD45 antibodies conjugated to three fluorophores, allowing us to test the seven different subjects in one well (A, B, C, AB, AC, BC, ABC). After barcoding, the samples were stained for CD56, CD3, CD14, Live/Dead, and IL-10, as in the above protocols. These cells were then pooled together and aliquoted into the Biolegend LEGEND screen plates to be stained with an individual PE antibody in each well. The cells were then then acquired on a CytoFLEX flow cytometer (Beckman Coulter) and analyzed via FlowJo 10.8.1.

Of the 350 antibodies in the LEGENDScreen*^TM^*, gMFI expression and %PE of 134 protein markers were productively stained on the NK cells. Two-tailed, paired T-tests were performed to determine significant differences between IL-10+ and IL-10-groups. The 134 protein p-values were adjusted in R using the Benjamini-Hochberg procedure with the p.adjust() function in R. Proteins were deemed significant if the adjusted p-value < 0.05. The gMFI and %PE values for significant proteins were placed into separate data.frames in R. gMFI values were mean-centered and scaled using the scale() function. The gMFI data were reshaped (reshape R package) using melt() to reformat for ggplot2 visualization before %PE values were appended. Final dataframe was plotted using ggplot() as a dotplot. The dot color represents the protein expression (gMFI) and the dot size represents the percent protein expression (%PE) for the significant proteins.

### Statistical analysis

Statistical significance is indicated by P values in figure legends and was determined by using Prism Versions 9 and 10 (GraphPad). The number of samples (n) and statistical comparison groups are indicated in figures and legends.

### Data availability

scRNA-sequencing data sets have been deposited in the Gene Expression Omnibus (GEO) under the following study accession: GSE266970.

### Human Study Approval

Informed consent was obtained from all participants. The institutional review boards of the National Institutes of Health and the University of Minnesota reviewed and approved this study.

## AUTHOR CONTRIBUTIONS

Designed research studies: S.E.H., G.T.H.

Conducted experiments: S.A.M., J.K.D., M.H.C., J.A.S., M.P.

Acquired data: S.A.M., J.K.D., M.H.C., J.A.S., M.P., K.S.B.

Analyzed data: S.A.M., M.M., J.K.D., M.P., G.T.H.

Provided samples: P.D.C., K.B.S.

Writing the manuscript: S.A.M., S.E.H., G.T.H.

Edited the manuscript: S.A.M., J.K.D., M.H.C., M.P., K.S.B, M.M., P.D.C, K.B.S, S.E.H., G.T.H.

S.A.M. and J.K.D. are co-first authors; The order reflects the relative contribution to the original manuscript.

## ACKNOWLEDGEMENTS

We gratefully acknowledge the following support services at the University of Minnesota that contributed to the research results reported: University Flow Cytometry Resource, University of Minnesota Genomics Center, and the Minnesota Supercomputing Institute. This work was supported by the following funding sources: NIH/NIAID R01AI143828 (S.E.H.), NIH/NIAID R01AI146031 (G.T.H.), NIH/NIAID T32AI007313 (J.K.D).

