## Supplemental Figures for "Phenotype and function of IL-10 producing NK cells in individuals with malaria experience"

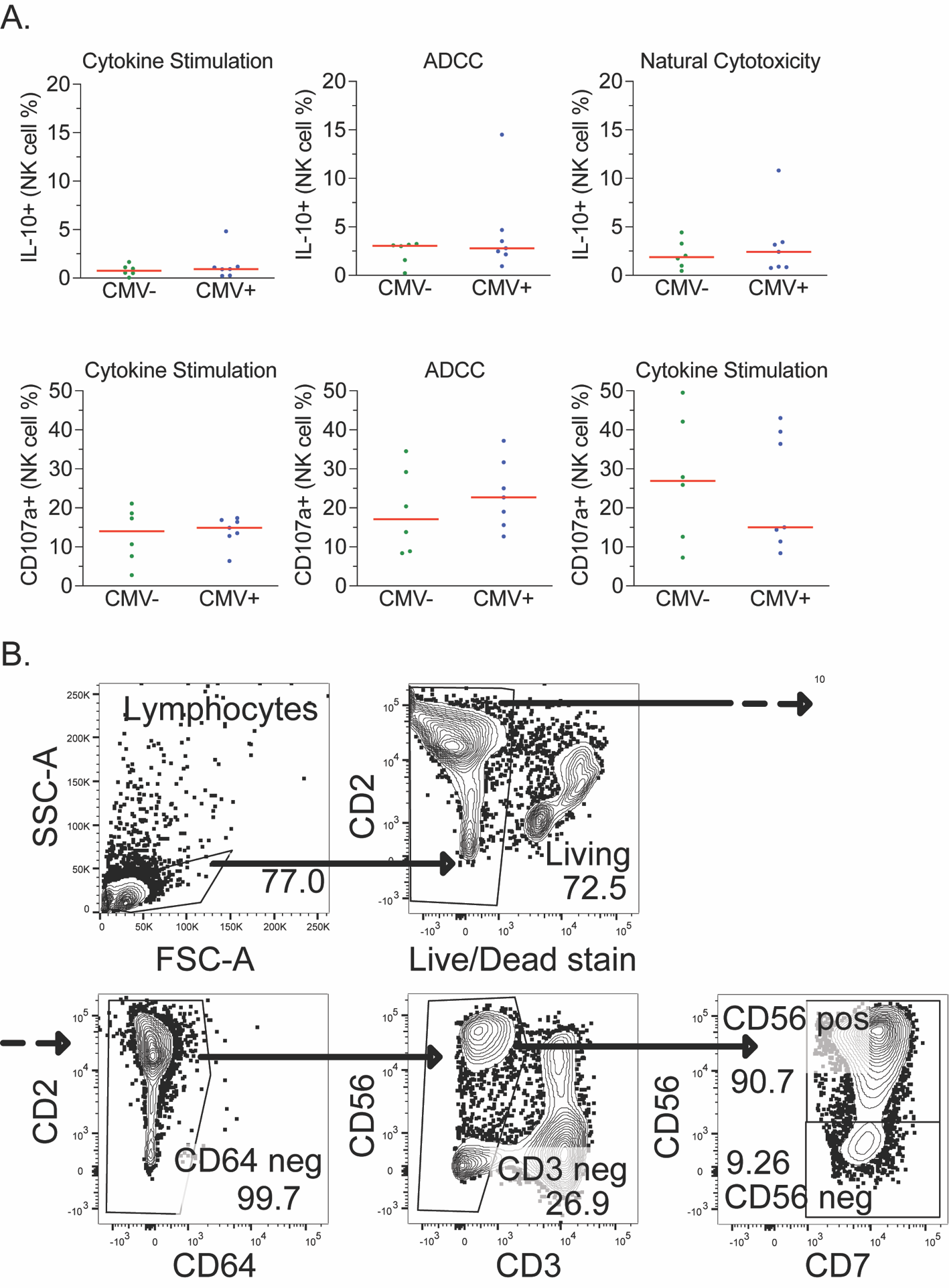

**Supplemental Figure 1. Analysis of CMV seropositivity in subjects and example Flow Cytometry gating. (A)** IL-10 (top row) and CD107a (bottom row) expression in CMV seropositive and seronegative individuals (USA). Groups were compared using Wilcoxon signed-rank tests (*p<0.05, **p<0.01, ***p<0.001); orange bars represent median values. (B) Example gating of a representative malaria experienced subject’s cells via flow cytometry.

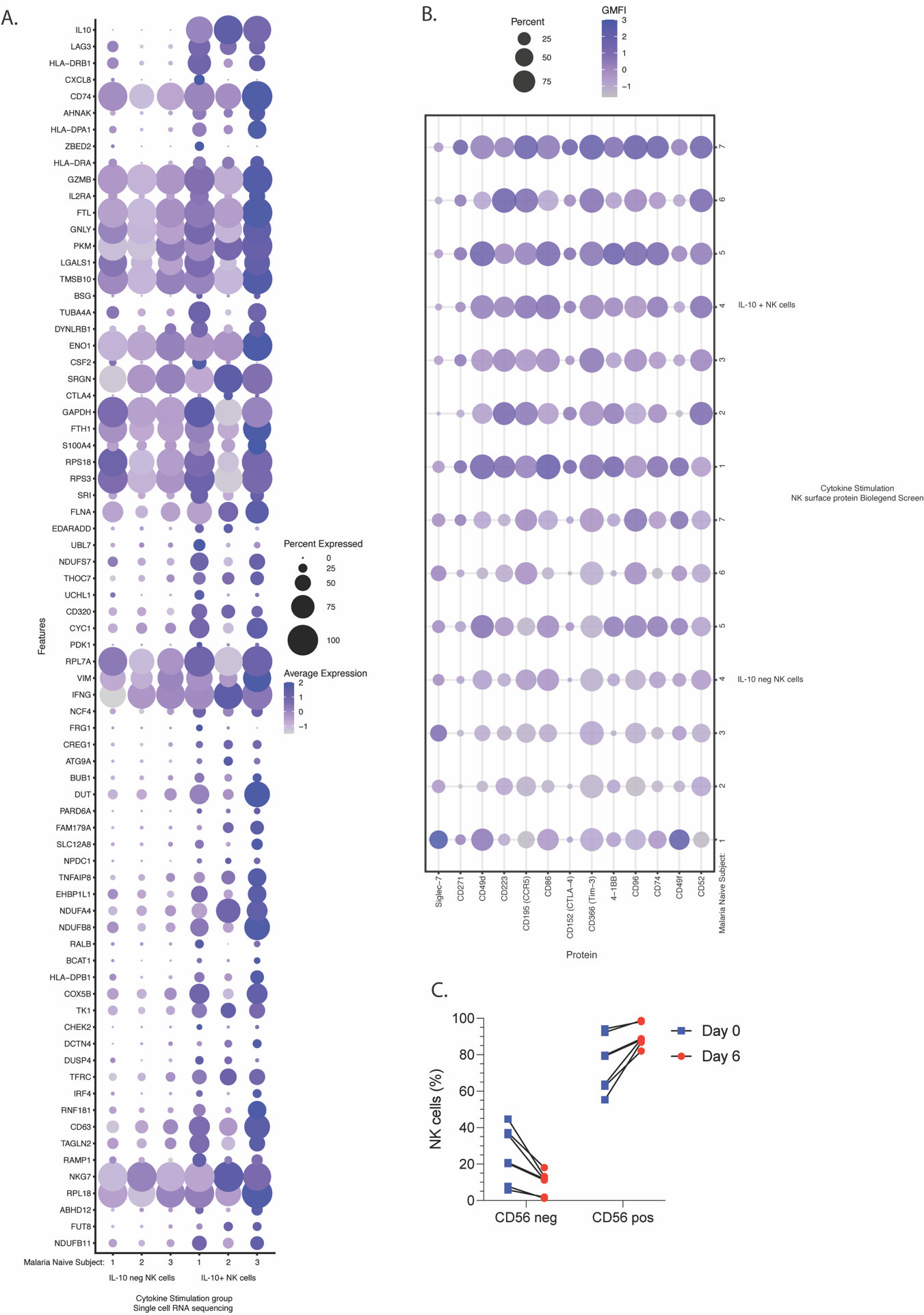

**Supplemental Figure 2. Initial single cell RNA sequencing and surface protein screen to identify potential additional markers associated with IL-10 producing NK cells.** (A) Single-cell RNA sequencing was done on three malaria-naïve individuals NK cells that had been through the Cytokine stimulation protocol. In the identified NK cells, IL-10 negative and IL-10 positive NK cells were compared, and the top significant candidate markers are shown. Circle size indicates percent of NK cells positive for the gene. Color intensity indicates expression level up or down. (B) Using a the Biolegend LEGENDSceen surface receptor screen, six malaria-naïve individuals NK cells that had been through the Cytokine stimulation protocol were test for surface expression of 350 proteins. IL-10 negative and IL-10 positive NK cells were compared, and significant candidate markers are shown. Circle size indicates percent of NK cells positive for the gene. Color intensity indicates expression level up or down. Data was analyzed using students t-test with Benjamini-Hochberg adjusted p-values. (C) Proportion of CD56 positive and CD56 neg that are Living, CD64-, CD3-, CD7+ at Day 0 and Day 6 post cytokine incubation.

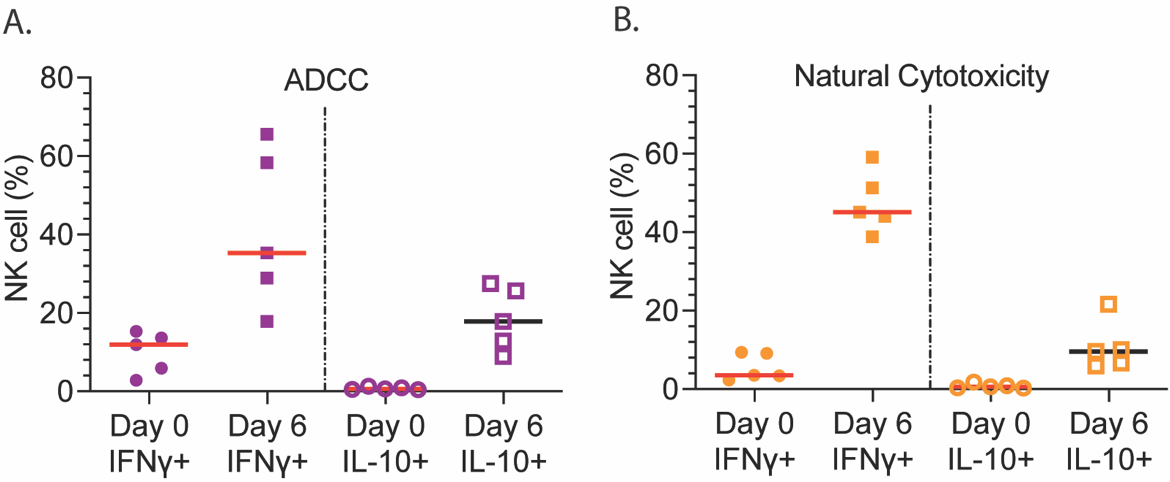

**Supplemental Figure 3. NK cell production of IFNγ or IL-10 directly *ex-vivo* (Day 0) or 6 days after stimulation.** (A-B) Malaria-experienced NK cells producing either IFNγ (Left) or IL-10 (Right) on Day 0 or 6 during ADCC (A) or Natural Cytotoxicity (B).

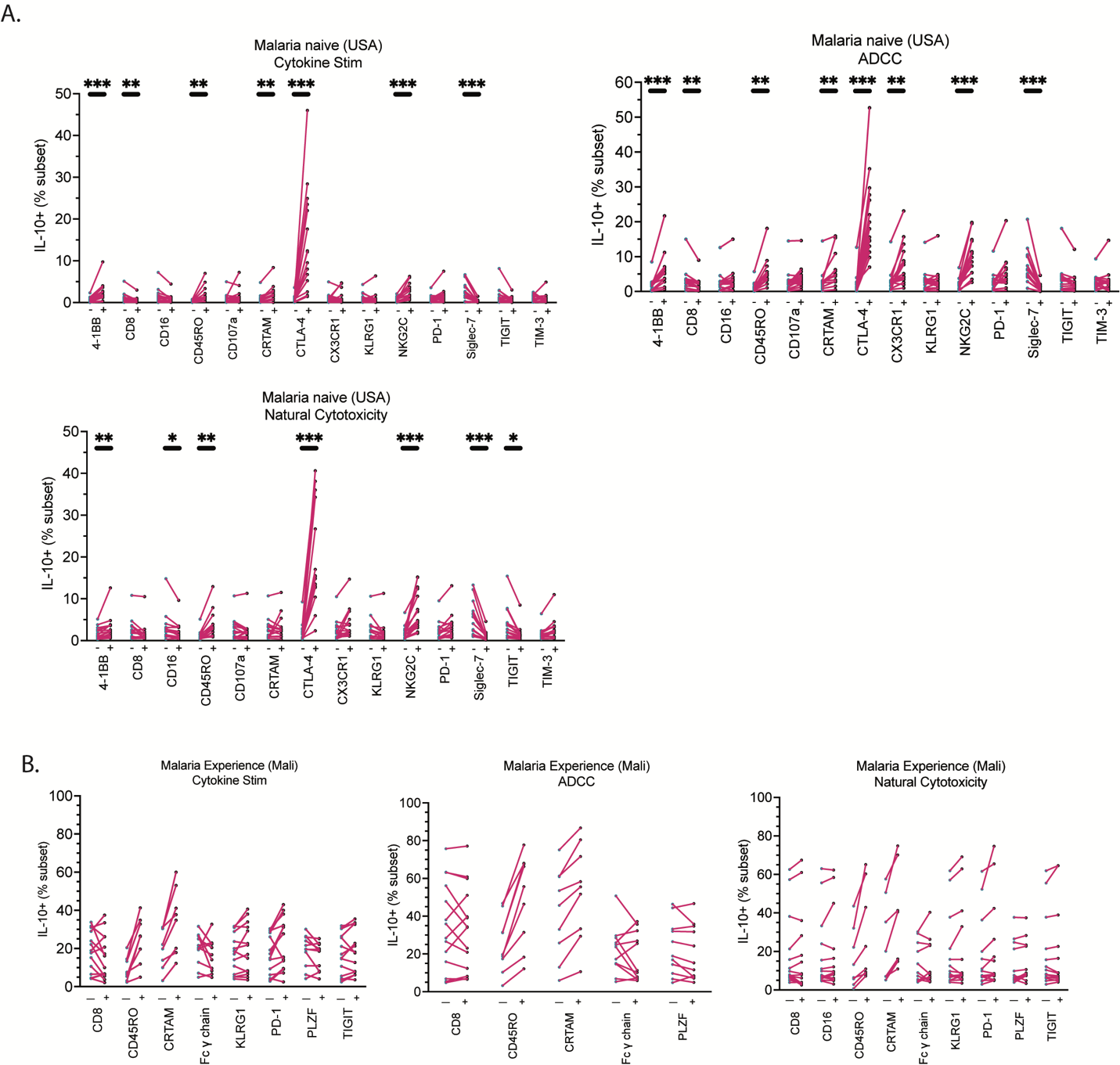

**Supplemental Figure 4. IL-10 production delineated by NK cell marker for malaria- naïve (USA) and malaria-experienced (Mali) individuals.** (A-B) IL-10 production by NK cells for Cytokine stimulation, ADCC, and Natural Cytotoxicity. Malaria-naïve (A) and malaria-experienced (B) IL-10 production by NK cells. Mann-Whitney non-parametric paired test done initially followed by a Bonferroni correction (14) to get the final p-value. * = p-value <0.05, ** = p-value <0.01, *** = p-value <0.001, **** = p-value <0.0001 post Bonferroni correction.

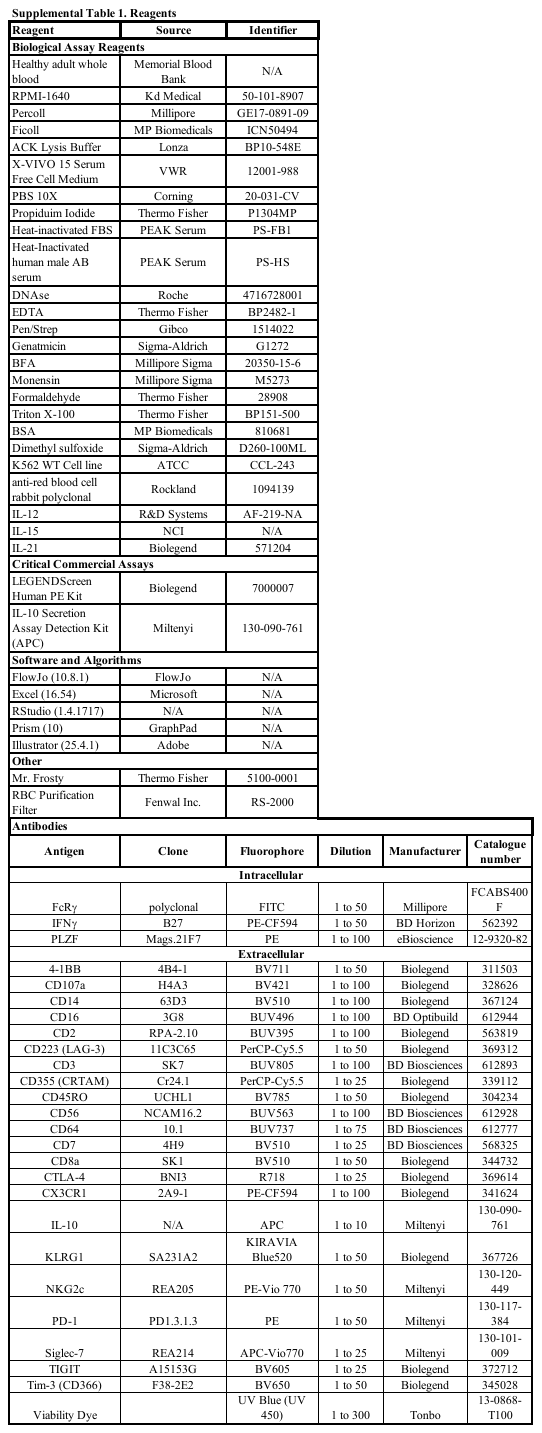
